# Prioritization of disease genes from GWAS using ensemble based positive-unlabeled learning

**DOI:** 10.1101/2020.07.12.199273

**Authors:** Nikita Kolosov, Mark J. Daly, Mykyta Artomov

**Affiliations:** ITMO University, St. Petersburg, Russia; Analytic and Translational Genetics Unit, Massachusetts General Hospital, Boston, USA; Broad Institute, Cambridge, USA; Institute for Molecular Medicine Finland (FIMM), Helsinki, Finland

## Abstract

Major complication in understanding disease biology from GWAS arises from inability to identify a complete set of causal genes. Integration of multiple omics data sources could provide an important functional link between associated variants and candidate genes. Machine-learning could take advantage of this variety of data and provide a solution for prioritization of disease genes. Yet, classical positive-negative classifiers impose strong limitations on the gene prioritization procedure, such as lack of reliable non-causal genes for training.

Here, we developed a novel gene prioritization tool - Gene Prioritizer (GPrior). It is an ensemble of five positive-unlabeled bagging classifiers, that treat all genes of unknown relevance as an unlabeled set. GPrior selects an optimal combination of algorithms to tune the model for each specific phenotype.

Altogether, GPrior fills an important niche of methods for GWAS data post-processing, significantly improving the ability to pinpoint disease genes compared to existing solutions.

## Introduction

Despite the tens of thousands of genetic associations identified using GWAS to date, the ultimate goal - informing and guiding therapeutic development - has been achieved for only a few phenotypes. A major complication in understanding disease biology from GWAS often arises from inability to directly identify disease genes ^1^. Therefore, additional post-GWAS analysis is needed to first, identify a variant that drives the signal within the locus, and then to connect this variant to a gene.

Fine-mapping, based on a Bayesian framework, sets out to prioritize variants within the locus and, ultimately, identify the disease-causing variant ^2,3,4^. Fine-mapping algorithms - FINEMAP ^5^, PAINTOR ^6^, fGWAS ^7^, SUSIE ^8^ etc. had significant impact on the field and had successfully identified causal variants for multiple traits. Importantly, fine-mapping is done independently for each locus and in its current configuration does not take advantage of biological relatedness (e.g., same pathway membership) of genes involved in a phenotype ^9^.

At the same time, identification of the disease gene linked to a disease-causing variant presents a major and yet unresolved challenge. Most GWAS associations implicate a set of correlated genetic variants, none of which alter the protein-coding sequence of a gene and which often physically span or are near to multiple genes. Since our knowledge of regulatory sequence patterns of the genome, the relevant cells, tissues and developmental time points relevant to disease are all incomplete, it is currently the case that the vast majority of GWAS ‘hits’ do not have a certain link to a gene. Though data sets with which to infer functional annotation and gene expression are growing rapidly in their utility.

Post-processing of the GWAS results with inclusion of functional information is a promising direction on the road to identify disease genes. For example, Post-GWAS Analysis Platform (POSTGAP^10^) uses GWAS summary statistics along with LD-structure and external functional databases (GTEx ^11^, FANTOM5 ^12^, RegulomeDB ^13^ etc.) to prioritize SNPs within the locus and narrow down the list of potential gene candidates. Yet, the gene prioritization utility of POSTGAP is still in early development.

Altogether, fine-mapping, functional annotations and known biologic relatedness across putative disease genes become valuable data sources for gene prioritization, defined as evaluation of the likelihood of gene involvement in generating a disease phenotype ^14^. Machine-learning (ML) based prioritization could take an advantage of these data sources and provide a solution for novel disease gene identification.

Typically, existing ML solutions use Positive-Negative (PN) classification strategy. In PN classification per-gene probabilities are obtained by using known disease genes as a positive (P) training set and unknown genes as a negative (N) training set ^15, 16, 17, 18^. Such an approach suffers from contamination of a negative set by hidden positives (HP), represented by yet undiscovered disease genes. Additionally, it is challenging to find reliable negative examples (i.e., genes that with certainty do not contribute to the development of a phenotype). Most biological databases do not store negative evidence (e.g., absence of gene interaction), rather they provide only observed positive evidence. As a result, PN-classifiers could suffer from high false-negative prediction rates and biased quality metrics.

It is feasible to design a model, where a limited number of reliable positive examples (likely causal genes) will be used along with the rest of genes without treating the latter as reliably negatives. Positive-Unlabeled (PU) learning has been developed to overcome limitations of PN-learning. PU-learning treats unknown examples as a mixture of P and N, called unlabeled (U) set.

PU-learning was first proposed by Denis et al. ^21^, and several algorithms have been published since then ^19,20,21^. A particular class of PU methods - PU-bagging, showed the best stability of the learning algorithm ^22^. Specifically, Mordlet et al. ^22^ proposed the “bagging SVM” approach that took advantage of a limited number of positive examples and significantly improved performance and stability of classification using a new aggregation technique.

Nevertheless, a single ML algorithm cannot fit all complex phenotypes and highly heterogeneous biological data. To overcome this, Yang et al. ^23^ introduced a concept of integration for several PU learning classifiers into one workflow using ensemble technique. This technique was only tested with a specific family of PU algorithms - two-step methods, heuristic in nature and sensitive to the initial choice of negative examples ^24^, significantly limiting applicability to GWAS data. Two-step PU algorithms first attempted to identify negative examples in the unlabeled set, and then train a model from the positive, unlabeled and likely negative examples. However, directly learning to discriminate P from U with estimation of optimal misclassification costs leads to better results ^25^.

Therefore, combination of different ML algorithms (kernel-based and tree-based) along with PU bagging is a promising strategy for building a gene-prioritization model suitable for a large number of complex phenotypes and high variety of data sources, that is still lacking in the field.

Here, we propose a novel PU-learning based gene prioritization tool - Gene Prioritizer (GPrior), intended for post fine-mapping usage of GWAS results. In GPrior we implemented the ensemble of 5 different ML classifiers for PU-bagging with further selection of the optimal combination of predictions. Our approach returns probability scores for the whole provided set of genes based on similarity level with positive examples and is complementary to any other gene prioritization tools and fine-mapping techniques.

We illustrate the utility of PU-learning and GPrior with common public ML quality evaluation dataset with known ground truth and a series of case studies. Comparison with popular methods (TOPPGENE ^26^; Bagging SVM ^22^; MAGMA ^27^) confirmed significantly higher quality of predictions returned by GPrior.

## Methods

GPrior was designed for prioritizing disease-relevant genes given a matrix of gene-level features and a set of reliably causal genes. We integrated multiple ML techniques in a single tool and a data-driven framework to select the most appropriate algorithm (or combination of algorithms) on a case-by-case basis.

The prioritization scheme includes two independent steps. First, each ML algorithm is used for positive-unlabeled bagging and generation of predictions for each gene. Second, the best-performing combination of predictions is generated. To ensure the independence of steps, a set of true genes that is used for training is separated into two parts – set of genes for training individual ML algorithms and an algorithm evaluation set (**Figure 1**, **Sup. Figure S1**). The latter is used to evaluate the quality of predictions from ML algorithms and select predictions that will contribute to the optimal combination. Altogether, such an approach allows to combine multiple learning algorithms and achieve previously unattainable for an individual algorithm performance.

**Figure 1.**
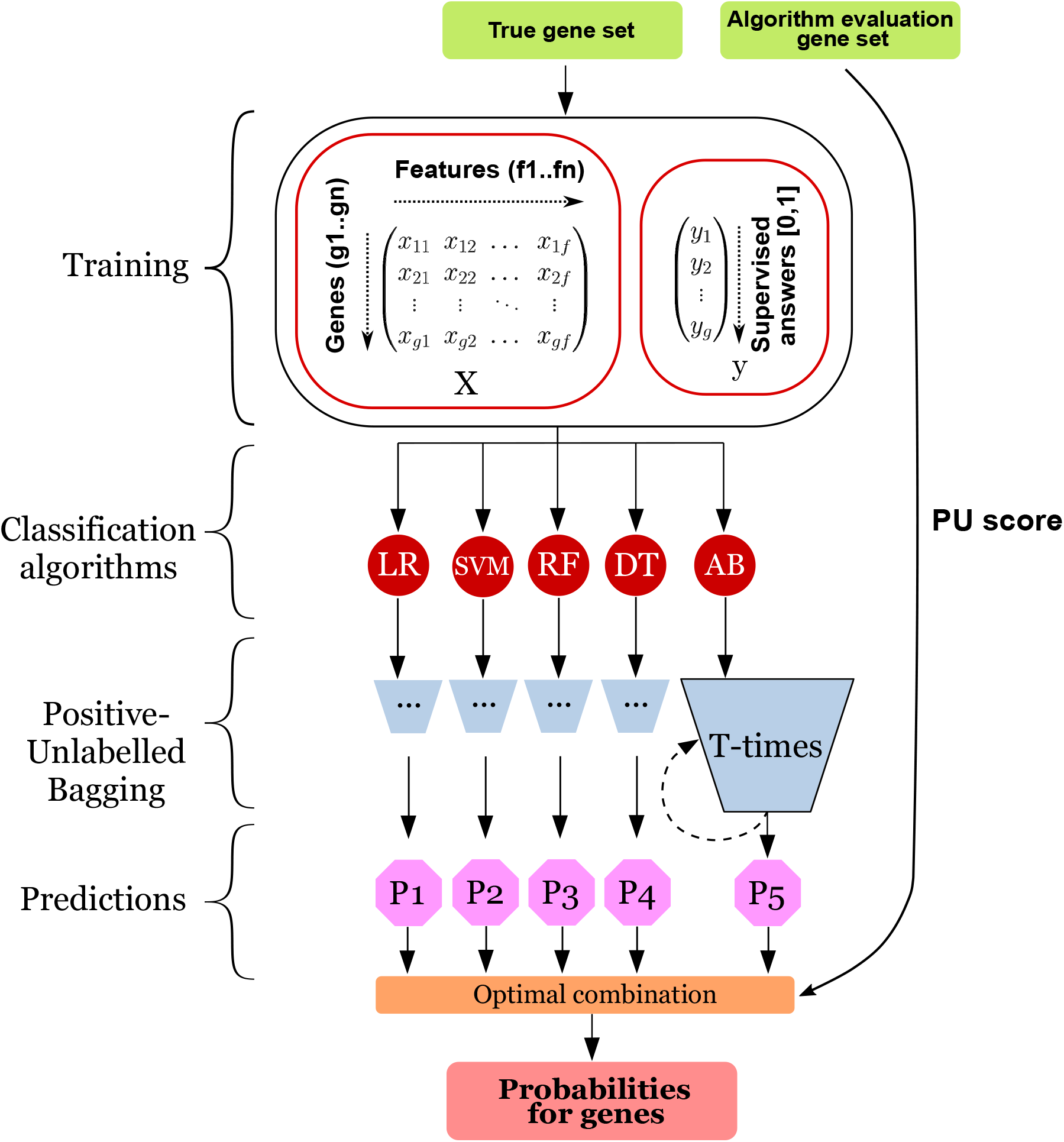
GPrior ensemble positive-unlabeled learning framework. Matrix of gene features along with vector of supervised answers is used to train 5 models using PU-bagging approach. Two independent gene sets are used for training – true set of genes for individual classification algorithms training and algorithm evaluation set of true genes for selecting the optimal combination of the predictions. Predictions are generated using positive-unlabeled bagging and further an optimal combination returning the largest *PU*-score is returned.

### Input and Features

In addition to the described above true gene sets needed for training, GPrior requires a data matrix with rows representing genes and columns representing features.

GWAS summary statistics contains only variant information which needs to be converted into gene level data. Initially, we filtered out likely non-associated variants with the threshold determined on case-by-case basis, ensuring inclusion of the majority of potentially causal genes (even if no significant association was observed in GWAS) into the prioritization analysis. Otherwise, GWAS *p*-values were not a part of the prioritization model.

Next, we used POSTGAP ^10^, which takes advantage of LD structure and variant functional annotations to assign potential gene candidates for each variant. Such preprocessing of GWAS summary statistics yields a variant-based data matrix with mappings to a list of candidate genes.

A major challenge in transforming variant-based data into gene-based data matrix for GPrior is preservation of valuable information about variant annotations. We used a transformation of variant-level features (e.g. functional annotations, GERP scores, etc) into gene-level features using a method proposed by Lehne et al. ^28^ to obtain a gene-based data matrix.

In addition, we used gene expression and gene interaction data that proved their utility for the gene identification problem in previous works ^15, 29, 30^. Specifically, we used the GTEx database to obtain median gene expression levels for 53 tissues and Reactome ^31^ and UCSC (GeneSorter) ^32^ databases were used for gene interaction data (**Sup. Table S1**).

Additional functional features and predictions of other prioritization algorithms could be included in the data matrix to be used for GPrior model to boost the performance quality ^33^. GPrior could take as an input either the raw output of POSTGAP (variant-based data matrix) or any gene-based data matrix provided by a user.

We kept the same set of features for the case studies to preserve the fairness of performance comparisons for different phenotypes. Although, for each phenotype features could be selected in concordance with phenotype-specific needs, for example, relevant cell type expression data. Overall, GPrior is not bound to a pre-specified set of features and could be used with any user-defined set of features to boost the trait-specific performance.

### Feature preprocessing

As for any ML approach, prior to algorithm execution, features should undergo preprocessing procedure to equalize scales and eliminate potential performance biases.

In gene prioritization, raw data sets can potentially reach hundreds or thousands of features. Along with a limited number of positive examples, this enormity can lead to the “curse of dimensionality” ^34^. Hypothetically, the more features are available in the data the more accurate result should be expected. However, greater number of features leads to the exponential growth of the training examples amount, required to cover the sparse feature space and achieve acceptable prediction quality. In real-life applications, the number of positive examples is limited, therefore, conventionally this problem is solved by clustering raw features. GPrior uses agglomerative feature clustering as a dimensionality reduction technique to extract appropriate number of features and achieve the highest performance (**Suppl.Methods, Input and Features**).

### GPrior algorithm

GPrior consists of five PU Bagging ensembles, each of them uses a different classification algorithm: Logistic Regression (LR), Support-Vector Machine (SVM), Decision Tree (DT), Random Forest (RF), Adaptive boosting (AB) (**Figure 1, Sup. Methods, GPrior Algorithm**).

Each positive-unlabeled bagging procedure starts with a creation of a training set with all positive (P) instances, treated as Positives, and a random subsample of unlabeled, size of P, treated as negatives. Resulting in the size of a bootstrap sample being equal to P. This way, on each iteration only a small portion of unlabeled instances is treated as negatives, minimizing false negative error rate (**Sup.Figure S2**). Each learning method is then fine-tuned by finding an optimal set of parameters (**Sup. Table S2**).

After training and tuning, individual classifiers generate a probability score for each gene to belong to the positive class. All the steps are repeated *T* times. Per-gene probabilities are obtained by dividing the sum of all predictions by the number of times each gene was sampled from the unlabeled set. All the predictions are averaged and stored as a final PU Bagging result. All the steps are repeated for each classification algorithm.

Next, GPrior selects the combination of predictions that shows the best performance in prioritizing true genes from an independent algorithm evaluation set. Since “true negative” data points that falsely were classified as positives could not be identified in PU-data, any metric depending on false positives could not be applied for quality evaluation. We used *PU-*score as a formal quality metric suitable for positive-unlabeled data classes ^25, 35^ (**Sup. Methods**).

All combinations from individual predictions are evaluated using PU-score calculated for algorithm evaluation set and the best performing composition of methods is then selected as the best fitting for a given phenotype. Selected combination is used to return a vector of probabilities corresponding to the genes in the input matrix.

## Results

### Performance evaluation

The number of known true positive and negative data points is a critical information parameter for gene prioritization. It is challenging to estimate both the number of genes involved in a complex trait and the number of genes confidently irrelevant for the disease. Height, as a classic example of a highly polygenic trait, shows effect size for a median GWAS SNP in the genome about 10% of that for genome-wide significant hits. This suggests that in current GWAS for height, significant associations are observed only for a small proportion of true positive data points, while many others are yet to be confirmed. However, additional alleles at known genes are a likely source of much of what is missing so this does not easily translate into an estimate of how many relevant genes are implicated from a GWAS. As shown in recent works, the genetic architecture of height is broadly similar to that of a wide variety of other quantitative traits and diseases ranging from diabetes or autoimmune diseases to BMI or cholesterol levels ^36^ – for all of which the evidence suggests many more positive genes exist in the ‘currently not associated’ gene set. Additionally, only some of the genome-wide significant loci were mapped to a single gene, even further reducing the number of known true genes suitable for training a model. Therefore, it is reasonable to assume that gene prioritization algorithms should expect to be trained using only a small fraction of all disease-relevant genes.

Due to inability to obtain a GWAS dataset with confidently identified complete sets of the disease genes and disease irrelevant genes, for initial GPrior performance evaluation we used a public benchmark dataset. Breast Cancer Wisconsin (Diagnostic) Data Set - a popular public dataset for machine learning tools benchmarking was used to compare performance of GPrior to conventional PN-learning (biased SVM ^37^) and single PU-learning algorithm - bagging-SVM ^22^ (**Suppl.Methods, GPrior Performance Evaluation**). The data with known true positives and true negatives enables calculation of fair prediction error rates which is impossible for real life data with yet undiscovered true data points. While this dataset is not related to the gene prioritization problem, it clearly illustrates the benefits of ensemble PU-learning in case of only a small fraction of known true instances to be used for training.

The dataset contained 569 samples and 30 features (**Figure 2A**). Initially, we assumed that only 4% of true positive class samples (malignant tumors in the Breast Cancer Wisconsin data), are known and available for training. Since GPrior requires two independent true sets for training, the original known positive class was broken down into two parts – 5 samples were used for training and 3 for algorithm evaluation set. For all other methods the whole known positive class was used as a single batch.

**Figure 2.**
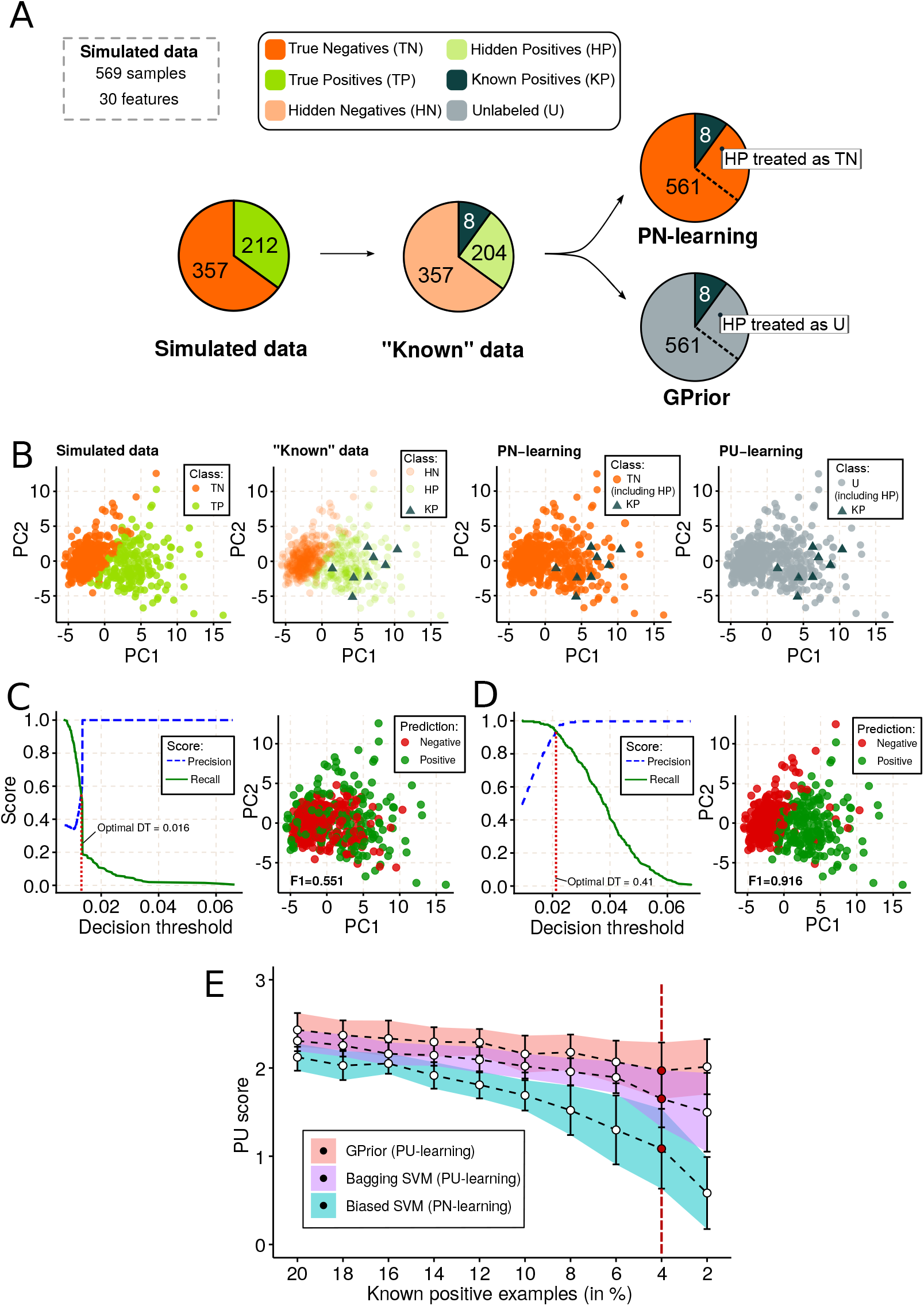
Benchmark dataset shows advantage of PU learning over PN learning in a scenario when few positive instances used for training. **(A)** Dataset breakdown. Known data represents scenario when only a part of the true positive instances were discovered to date; **(B)** PCA of the simulated data with highlighted true instance classes; **(C)** Prediction results for PN-learning (biased SVM) approach; **(D)** Prediction results for GPrior; **(E)** Performance of PU and PN learning approaches with respect to a fraction of known positive data points.

We performed PCA to highlight how data classes are recognized in PN and PU-learning (**Figure 2B**). PN-learning treats all instances as true negative, except those used for training (known positives). This way, PN-learning attempts to identify samples falsely classified to be true negative. In opposite, PU-learning treats all instances not used for training as unlabeled and is, therefore, free from assumption that true negative class exists. GPrior generated predictions using each algorithm and algorithm evaluation set was used to estimate *PU*-scores (**Suppl. Figure S3A**). All combinations of 5 algorithms available in GPrior were tested and the combination with the best *PU*-score estimated from the algorithm evaluation set was selected to return the results (**Suppl. Figure S3B,C**).

Precision-recall analysis was used to determine optimal decision threshold for each method and ensure that best possible performance was extracted from each algorithm (**Figure 2C,D**). As a result, GPrior carefully identified corresponding sample classes (F1 = 0.916, **Suppl.Figure S4A, B**), while PN-learning fails to achieve similar performance (F1=0.551, **Suppl.Figure S4A, C**).

Testing all 3 methods - biased SVM, bagging SVM and GPrior for different fractions of known positives (100 simulations) results in superior performance of the latter (**Figure 2E**). Notably, PN-learning starts to behave equally well compared to PU-learning only if more than 30% of the true positive data points are already known and used for training.

For the benchmark dataset, with pre-specified true positives it is possible to compare *F1*-score and *PU*-score metrics. Both metrics show similar results, justifying further usage of *PU*-score for GPrior performance evaluation in case studies, where *F1*-score could not be computed (**Figure S4D, E**).

Additionally, we tried to fix contamination of unlabeled-class with hidden positives and change only the number of KP without changing the percentage of contamination to make the setup even more fair for PN and PU learning. Despite this, we obtained better results from GPrior (**Figure S5**).

To illustrate utility and performance quality of GPrior we performed several case studies using GWAS results for several phenotypes: inflammatory bowel disease (IBD), educational attainment (EA), coronary artery disease (CAD) and schizophrenia (SCZ).

### Case study 1: Inflammatory bowel disease (IBD)

We used GPrior and summary statistics from Huang et al. ^38^, to construct gene prioritization for IBD. Summary statistics was preprocessed to obtain a data matrix with 2,166 gene candidates found in loci with original p-value < 10^−8^. A list of 31 genes with known evidence to be likely causal for IBD was used as a positive training set. Algorithm evaluation set consisted of 14 genes reported in monogenic loci with *p*-value < 10^−10^ found in GWAS catalog. Independent validation set used only for performance evaluation included 51 genes found within monogenic loci with *p*-values falling in range 10^−10^ - 10^−8^ (**Figure 3A, Suppl. Table S3**).

**Figure 3.**
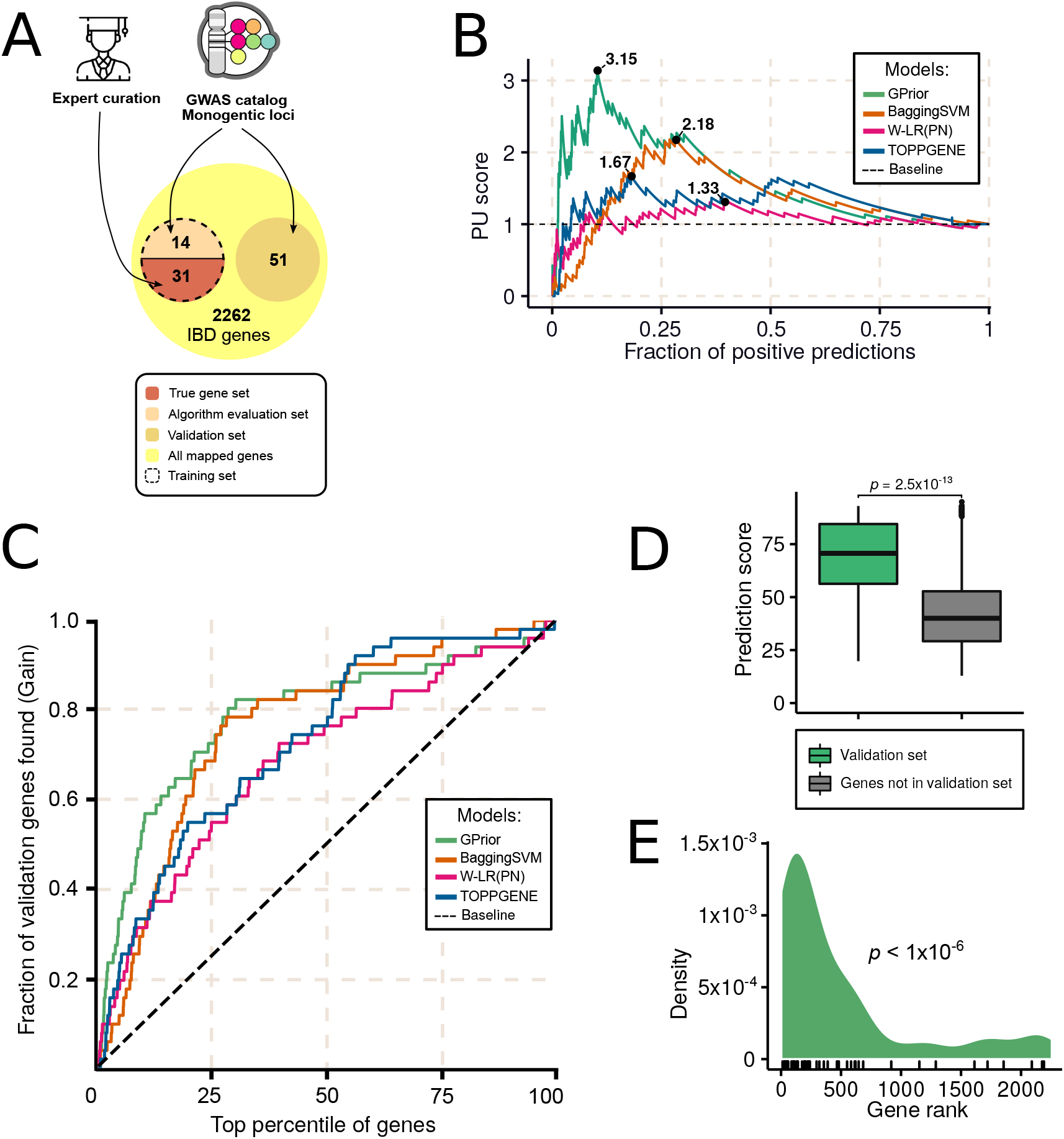
Gene prioritization for inflammatory bowel disease GWAS. **(A)** Scheme for selection of training, algorithm evaluation and validation gene sets; **(B)** Classification quality comparison for GPrior, Bagging SVM and conventional PN-learning with weighted linear regression; **(C)** Cumulative gains curve shows better prioritization of true genes at the top of the candidate list using GPrior in comparison with other methods; **(D)** True genes from the independent validation gene set receive significantly higher scores than genes found within the same loci but not implicated in the disease; **(E)** Enrichment of true genes from independent validation gene set among top predictions from GPrior.

We generated gene priorities using GPrior (**Sup. Table S4**) and a set of methods for comparison - a single PU-learning (bagging-SVM ^22^), PN-learning (weighted LR ^25^) and TOPPGENE ^26^. While GPrior implies two training steps and usage of two training gene sets – true gene set and algorithm evaluation set, for other methods we used a union of the two gene sets for training.

Next, we compared the performance quality of the methods. *PU*-score is a formal quality metric for a ML-based classifier, rather than for a prioritization itself, and it depends on the decision threshold used to assign classes to the instances. Gene prioritization implies only a construction of the ranked list of genes, but not the classification of the genes into “disease” and “non-disease”. Yet, we evaluated the maximal possible performance of the methods in the classification problem. We used an independent validation gene set to estimate *PU*-scores for all possible decision thresholds (fraction of positive predictions made by the classifier) and GPrior has significantly greater maximal *PU-*score compared to others (**Figure 3B**).

To evaluate prioritization quality, we estimated cumulative gains. Gain chart shows enrichment of the genes from the validation set at the top of the ranked list of predictions, that is the sharper is the growth of gain in the beginning of the chart – the more enriched are correct predictions at the top of the predictions list (**Figure 3C**).

Since original GWAS summary statistics was preprocessed to include only variants with *p-*value < 10^−8^, all 2,166 genes in the data matrix are found in or in a proximity of significantly associated loci. GPrior does not use association strength or DNA location information for gene prioritization. Yet, genes from the validation set are significantly prioritized over the non-relevant neighbors (Mann-Whitney, one-sided, *p*-value = 2.5×10^−13^, **Figure 3D**).

We evaluated non-randomness of the predictions, by estimating enrichment of the validation set genes at the top of the ranked list produced by GPrior (permutation p-value < 1*10^−6^; **Figure 3E**).

Treatment of all genes from the validation set as a finite set of disease genes, implies that all genes that are not in the validation set are true negatives. In case of polygenic traits, this is most likely a false assumption which would lead to an underestimation of the true value of the area under the *ROC*-curve (ROC AUC). Thus AUC values will illustrate only approximate quality measurement. In such settings, GPrior demonstrated the most efficient predictive power out of all tools (AUC = 0.8, **Suppl. Table S5**).

### Case study 2: Educational attainment

We performed a control experiment to demonstrate that GPrior predictions are disease specific and are driven by underlying biological similarities for disease related genes. We considered two phenotypes with likely very modest overlap in underlying biological causes – IBD and educational attainment (EA). We hypothesized that usage of the training gene set fitted for IBD should fail to predict genes for EA.

GWAS summary statistics from Lee et al ^39^ for EA was preprocessed to obtain the data matrix of candidate genes (N=10,638) and features (**Methods**). To eliminate potential bias in the size of training sets for the two phenotypes, we used for GPrior training only 18 genes (12 for ML training and 6 for algorithm selection) from the IBD training gene set that were also found in EA GWAS loci with *p*-value < 10^−6^. As a validation set for IBD we used the original IBD validation genes (N=51), for EA we used 381 genes found in monogenic loci from GWAS catalog EA results (**Sup. Table S6, Sup. Methods**).

Usage of appropriate training set for IBD resulted in significant enrichment (permutation P<10^−6^) of validation set genes in the top predicted genes (**Sup. Figure S6A**). Predictions based on the same list of training genes were constructed for EA and demonstrated no enrichment of the EA-specific validation gene set (permutation *p*-value p-value =0.12, **Sup. Figure S6B**). Yet, usage of the EA-specific training gene set of the same size (N=18, **Sup.Methods**) led to successful prioritization of EA-specific genes (permutation P<10^−6^, **Sup. Figure S6C**).

We expanded the training set for EA by inclusion of all genes found in monogenic loci in GWAS Catalog with (N=119) and repeated prioritization analysis. As a result, we obtained even more significant enrichment for the validation set and confirmed superior performance quality of GPrior in comparison with other methods (**Sup. Figure S7, Sup. Tables S7,S8**).

Finally, we estimated how strongly initial GWAS summary statistics preprocessing contributed to the overall success of the prioritization. While POSTGAP, that was used for mapping variants to a list of candidate genes, is not designed specifically for gene prioritization, it reports a variant-to-gene mapping score based on the sum of values for 7 features (**Sup. Figure S8**). We used the maximal variant-to-gene score for each gene to construct a ranked list of genes. First, we estimated the largest possible *PU*-score for the model using only POSTGAP-based gene ranking for educational attainment data (*PU*-score = 3.82). We limited feature space to exactly the same 7 features and constructed gene prioritization using GPrior, resulting in ~10% increase in *PU*-score (*PU*-score = 4.1). GPrior uses all available features, while POSTGAP score is limited by only initial 7 features, yet expanding feature space for GPrior yields significant increase leading to the maximal *PU*-score of 4.84 (**Sup. Figure S7**), showing ~27% increase in quality.

### Case study 3: Coronary artery disease (CAD)

We used the summary statistics of coronary artery disease GWAS of 34,541 CAD cases and 261,984 controls from UK Biobank followed by replication in 88,192 cases and 162,544 controls ^40^. After preprocessing we obtained a gene-based data matrix with 2,794 gene candidates found in loci with original *p*-value < 10^−8^.

Recent review by Khera and Kathiresan ^41^ was used to compile gene sets for GPrior (**Figure 4A**). All genes with identified biological roles in any of the known disease pathways were used for the training set (TS=18, AES=8). All other genes, implicated in CAD, but with yet undiscovered molecular pathway membership became a validation set (VS=37) (**Suppl. Table S9**).

**Figure 4.**
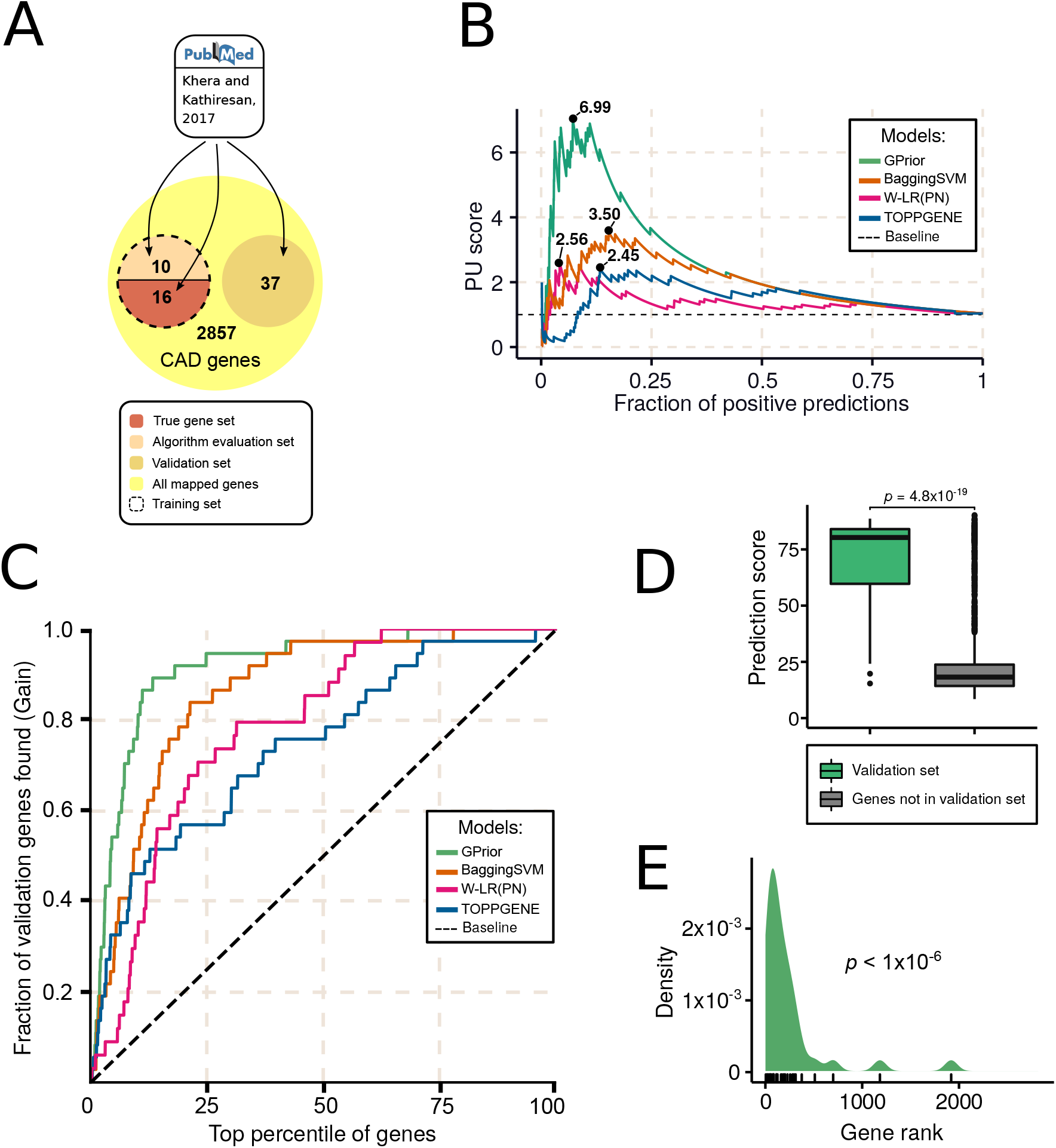
Gene prioritization for coronary artery disease GWAS. **(A)** Scheme for selection of training, algorithm evaluation and validation gene sets; **(B)** Classification quality comparison for GPrior, Bagging SVM and conventional PN-learning with weighted linear regression; **(C)** Cumulative gains curve shows better prioritization of true genes at the top of the candidate list using GPrior in comparison with other methods; **(D)** True genes from the independent validation gene set receive significantly higher scores than genes found within the same loci but not implicated in the disease; **(E)** Enrichment of true genes from independent validation gene set among top predictions from GPrior.

Prioritization list obtained with GPrior (**Sup. Table S10**) has shown the best accuracy with all quality metrics in comparison with other methods (**Figure 4B-E, Sup. Table 11**).

Conclusively, using risk genes with known molecular pathway membership GPrior successfully prioritizes genes with yet unknown biological contribution but confidently implicated in the disease. Importantly, by further analyzing feature importance in the prediction model it is possible to build testable biological hypotheses for novel genes discovered in predictions.

### Case study 4: Schizophrenia

We used GWAS Summary statistics from Pardiñas et al ^42^. This study used genotypes of 105 318 individuals, 40,675 schizophrenia cases and 64,643 controls.

After preprocessing we obtained a gene-based data matrix with 3,831 gene candidates found in loci with original *p*-value < 10^−6^.

Training set was prepared using reported genes found in monogenic loci from GWAS meta-analysis results ^42^. Training gene set for individual ML algorithms included 20 genes with *p*-values falling in range 10^−44^ - 10^−14^, and algorithm evaluation set included 24 genes with *p*-values within 10^−13^ - 10^−8^ range. Validation set (VS) included 28 genes and was obtained from the same study and included all genes from significant polygenic loci (**Figure 5A, Sup. Table S12, S13**).

**Figure 5.**
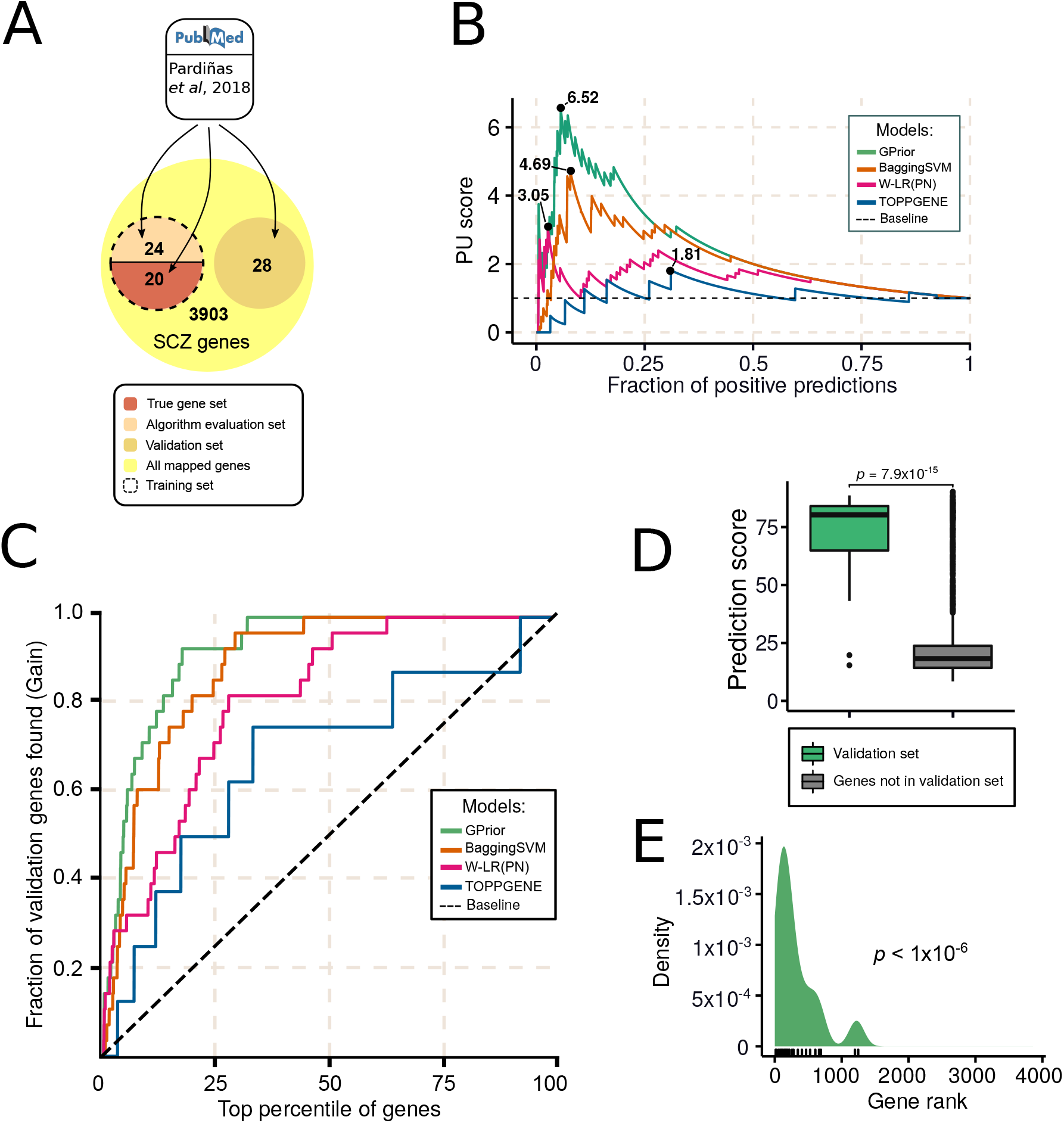
Gene prioritization for schizophrenia GWAS. **(A)** Scheme for selection of training, algorithm evaluation and validation gene sets; **(B)** Classification quality comparison for GPrior, Bagging SVM and conventional PN-learning with weighted linear regression; **(C)** Cumulative gains curve shows better prioritization of true genes at the top of the candidate list using GPrior in comparison with other methods; **(D)** True genes from the independent validation gene set receive significantly higher scores than genes found within the same loci but not implicated in the disease; **(E)** Enrichment of true genes from independent validation gene set among top predictions from GPrior.

GPrior demonstrated superior results in comparison with other methods using all quality metrics. GPrior achieved the highest PU score (9.64) and AUC (0.92) values. On all the top intervals of the predictions list (1%,5%,15%,25%) GPrior showed the highest enrichment of the validation set genes (**Figure 5B-E, Sup. Table S14**).

Conclusively, even for complex phenotypes with limited biological mechanism knowledge, like schizophrenia, GPrior is well powered to detect the relevant signal of biologic relatedness and prioritize likely disease genes.

### MAGMA comparison

We compared GPrior with a commonly used method that takes GWAS summary statistics as an input and attempts to pinpoint likely disease genes - MAGMA ^27^. It computes gene-based *p*-value (mean association of SNPs in the gene, corrected for LD). We ran MAGMA with default parameters and compared performance quality with GPrior. As an output MAGMA returns a list of genes and corresponding *p*-values, which we used to sort the list for prioritization purposes.

One of the challenges for a non-biased comparison was the relatively small number of gene candidates in output from MAGMA. Therefore, we took the same as reported in MAGMA number of top genes from GPrior results to compare equal number of gene predictions.

GPrior demonstrated enrichment of top ranked predictions for validation sets for all phenotypes – EA (*p*-value=9×10^−3^), schizophrenia (*p*-value=7×10^−4^), CAD (*p*-value=3×10^−3^) and IBD (*p*-value=0.019). MAGMA produced significantly enriched predictions only for CAD (*p*-value=9×10^−3^) (**Suppl. Figure S9**).

Conclusively, GPrior demonstrated best performance out of all evaluated approaches for gene prioritization in multiple settings and for various phenotypes.

## 4. Discussion

A large number of GWAS studies performed to date provide an invaluable source of information for generating biological hypotheses for disease causes. The majority of these studies greatly benefited from fine-mapping that implicated a limited number of gene candidates. However, for highly polygenic phenotypes like schizophrenia, known genes represent only a tiny segment of the disease biology.

The challenge of mapping “variants to function” could greatly benefit from machine learning approaches. Especially, for those phenotypes for which a strictly genetic fine-mapping approach has had limited success in conclusively identifying risk genes. As we illustrate, conventional positive-negative machine learning approaches require a substantial fraction of already known disease genes to achieve sufficient prioritization quality for novel candidates. Additionally, it is nearly impossible at this point to confidently state that a gene is not involved in a disease, therefore, directly assuming “negative” examples for training is fated to include false negatives in a training set, further reducing prediction quality.

Instead, we provide a tool that uses positive-unlabeled learning and requires confidence in selection of only positive instances for training. Such genes are relatively easy to identify based on association significance, previously reported functional studies, etc. Importantly, PU-learning performs well even when the training set is quite small.

Additional challenge for a single-method-based solution is presented by phenotype complexity. Phenotypes may present significantly different genetic architectures or impose certain limitations on the set of available data sources; therefore, it is unlikely that a single technique will be suitable for gene prioritization in all of them. We provide a software package for gene prioritization – GPrior that takes advantage of the ensemble of PU-learning techniques. Such approach overcomes unresolved challenges of PN-learning and issues arising from phenotype complexity. In GPrior, two key steps of the model training: PU-classifiers training and selection of optimal classifiers combination are performed using two independent gene sets. The two-step strategy ensures independent quality assessment for all classifiers and unbiased selection of the optimal prioritization method, as well as delivering optimal prioritization results for the specific phenotype.

GPrior can be utilized with many sources of functional data. Data types used in our case studies – tissue expression levels, Reactome pathway data and others represent only a small part of possibilities. Each phenotype study would significantly benefit from inclusion of additional features, such as – single cell expression data, specific protein-protein interactions, gene conservation metrics (pLI, LOEUF) and others. In our study we have not selected features to be specific to each phenotype, therefore, users can expect to see even higher performance in case of thorough feature selection. Additionally, GPrior can be straightforwardly integrated with conventional fine-mapping tools. One of the limiting steps in our GWAS processing scheme was naïve selection of gene candidates from each locus. More sophisticated preprocessing of the raw GWAS summary statistics with methods such as SuSie or FINEMAP to improve variant-to-gene mapping could significantly aid variant-level to gene-level features transformation. Finally, we used a relatively conservative set of features for gene annotations, which could be significantly expanded with phenotype specific annotations.

Altogether, GPrior fills an important and currently underdeveloped niche of methods for GWAS data post-processing, significantly improving the ability to pinpoint disease genes compared to existing solutions.

## Code availability

https://github.com/faramer86/GPrior

## Acknowledgements

Authors would like to thank Dr. Alexey Sergushichev (ITMO University) and Dr. Maxim Artyomov (Washington University in St. Louis) for helpful discussions.

