## Supplementary Materials for "Prioritization of disease genes from GWAS using ensemble based positive-unlabeled learning"

### Supplementary Methods

Nikita Kolosov[1,3], Mark J. Daly[2,3,4,*], Mykyta Artomov[1,2,3,4,*]

[1] – ITMO University, St. Petersburg, Russia
[2] – Analytic and Translational Genetics Unit, Massachusetts General Hospital, Boston, USA
[3] – Broad Institute, Cambridge, USA
[4] – Institute for Molecular Medicine Finland (FIMM), Helsinki, Finland

**Authors declare no conflict of interests.**

**Training and validation gene sets** 3
**Input and Features** 5
*SNP-level to gene-level features transformation* 5
*Features preprocessing* 5
*Feature clustering* 6
**GPrior Algorithm** 7
*Training and individual ML predictions generation* 7
*Combining results from multiple ML algorithms* 8
*Prediction Quality Evaluation* 10
**GPrior Performance Evaluation** 12
*Wisconsin breast cancer dataset* 12
*GPrior performance comparisons* 12
**Case studies** 16
*Educational attainment* 16
**Comparison with MAGMA** 20

### Training and validation gene sets

To avoid sampling bias, we used three different compiling strategies for training and validation gene sets: expert curation, publication-based and GWAS Catalog mining (**Figure S1**).

Each approach started with analysis of the initial list of all already known associated genes (i.e. using OMIM, GWAS catalog etc.). True set of genes that further should be used for training has to be divided into two parts – training gene set (TS) and algorithm evaluation set (AES). PU-bagging performs well even with a very limited number of positive examples. Therefore, it is better to put the most confident true genes into TS, leaving genes with smaller evidence for AES. Examples for each approach are considered in case studies.

Validation gene set (VS) is not required for running GPrior and was used only for independent performance evaluation of the final set of predictions from all prioritization algorithms tested in this work. Expectedly, the need to keep a list of true genes for validation, artificially decreasing the size of the training dataset, leads to an underestimation of actual performance.

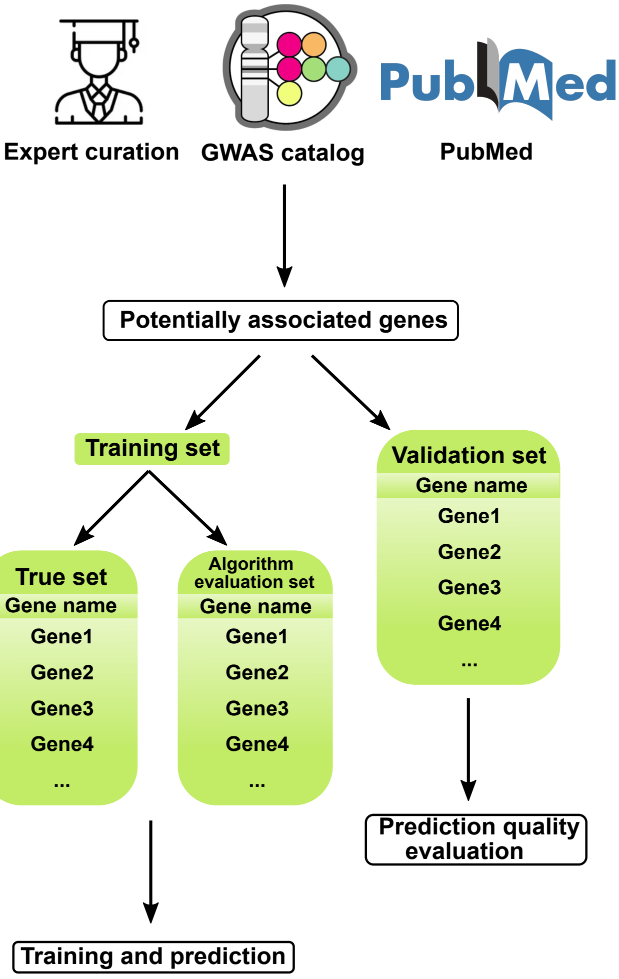


**Figure S1. Training and validation gene set selection**

We evaluated three approaches for true gene selection – expert curation, GWAS catalog and publication based. A list of selected likely causal genes is further binned into a validation set that is left aside and used only for evaluation of the final prediction quality in presented case studies and training set. The latter is split into a true gene set used for model training and algorithm evaluation set that is used for evaluating each of 5 PU-learning algorithms and selecting the optimal combination for final prediction.

### Input and Features

*SNP-level to gene-level features transformation*

We used Ensemble software (POSTGAP [1]) to pre-process GWAS summary statistics to link each variant to a set of potential gene candidates. POSTGAP considers all genes in the 1MB region from the variant and leaves only most likely candidates based on LD structure. Expectedly, many SNPs are mapped to the same candidate gene or, more frequently, a list of candidate genes.

SNP-level annotations, like variant functional consequences or GERP conservation scores of all variants linked to the same gene candidate should be summarized into gene-level features for further usage with gene prioritization algorithm.

Lehne et al. explored three approaches for integration of SNP-level features. First, *MaxT* method, selects the most extreme value for the SNP-level feature to be assigned to a gene (i.e. maximal GERP score across all SNPs mapped to a particular gene candidate). Second, *MeanT* computes arithmetic mean across all SNPs mapped to a gene candidate. Third, *TopQ*, computes arithmetic mean using only top quartile values of a SNP-level feature from the assigned SNPs.

Comparison of these three methods performed in *Lehne* et al. suggests that they have similar performance. We used both *MaxT* and *MeanT* to generate a redundant list of gene-level features. We excluded *TopQ* metric due to much higher computational complexity, but no expected quality advantage. As a result, each SNP-level feature was summarized for individual genes by computing maximum and mean values from a set of associated SNPs, resulting in two new gene-level features. Full list of gene-level features used for case studies is available in **Table S1**.

*Features preprocessing*

Standardized ML technique includes an important step of feature preprocessing prior to the model training. This is required to equilibrate feature scaling and eliminate technical biases in the data.

Feature scaling is used for normalizing the range of feature values. It is a common requirement for many distance-based machine-learning algorithms, including logistic regression (LR) and support-vector machine (SVM). It allows to compare differently scaled features and, more importantly, execute optimization techniques (i.e. gradient descent). We used a robust scaling method to scale all features. It subtracts the median from each feature value and scales the data according to interquartile range (IQR). IQR is the range between the 1st quartile (25th quantile) and the 3rd quartile (75th quantile). All of the features were processed as follows:

$$x' = \frac{x_i - Q_2}{Q_3 - Q_1}$$

where $x'$ – scaled feature value,

$x_i$ – original feature value,

$Q_{1,2,3}$ – feature distribution quartiles.

In comparison with standardization, where mean is subtracted from each feature value and all of the data is scaled according to variance, robust scaling is tolerant to outliers, since critical values are not so influential for IQR and median as for mean and variance.

*Feature clustering*

In gene prioritization initial data sets can reach hundreds or thousands of features. Along with a limited number of positive examples, this enormity can lead to the bias called curse of dimensionality [2]. Additionally, initial data sets usually contain some irrelevant or correlated features. Such biases can interfere with successful detection of hidden patterns within the data [3]. Together, unbalanced composition of number of features and positive examples degrades the performance of learning algorithms. Feature clustering is usually needed to determine appropriate number of features, corresponding to the number of positive examples available for training.

We used agglomerative feature clustering (FC). FC has been already used as an effective pre-processing method to enhance the discriminating ability [4, 5, 6, 7]. For the initial number of features defined as *n*, FC reduces the number of dimensions of initial dataset from *n* to *k* by grouping similar features together. During clustering features are merged based on the distance (e.g. Euclidean distance) between the clusters and selected linkage criterion (merging strategy). As a criterion we chose widely used Ward's method, defined as follows [8, 9]:

$$\Delta(A,B) = \frac{n_a \times n_b}{n_a + n_b} \times \|\vec{m}_a - \vec{m}_b\|^2,$$

$\Delta(A,B)$ - merging cost of combining the clusters A and B,

$n_i$ - is a number of points in cluster $i$,

$\vec{m}_i$ is a the centroid of cluster $i$.

The centroid is defined as the point that minimizes distance between itself and each point in the same cluster [9]. As a distance measure Ward's method uses Euclidean distance.

In other words, Ward's method produces a new cluster that has minimal merging cost. For feature clustering merging cost is calculated for all pairs of features. Two features with the lowest cost after merging are grouped together and averaged.

Unfortunately, the desired number of clusters *k* have to be defined in advance. Optimal *k* value is determined for each ML algorithm in GPrior by cross-validation.

### GPrior Algorithm

#### *Training and individual ML predictions generation*

GPrior consists of five PU Bagging ensembles, each of them uses a different classification algorithm: Logistic Regression (LR), Support-Vector Machine (SVM), Decision Tree (DT), Random Forest (RF), Adaptive boosting (AB).

Each positive-unlabeled bagging procedure starts with a creation of a training set with all *P* instances, treated as Positives, and a random subsample of *U*, size of *P*, treated as Negatives [10] (**Figure S2**). Resulting in the size of a bootstrap sample (*bN*) being equal to *P*. This way, on each iteration only a small portion of *U* is treated as *N*, minimizing false negative error rate. For each learning method in every iteration classifier was fine-tuned by finding an optimal set of hyperparameters, using a 3-fold cross-validation over a grid of hyperparameter tuples.

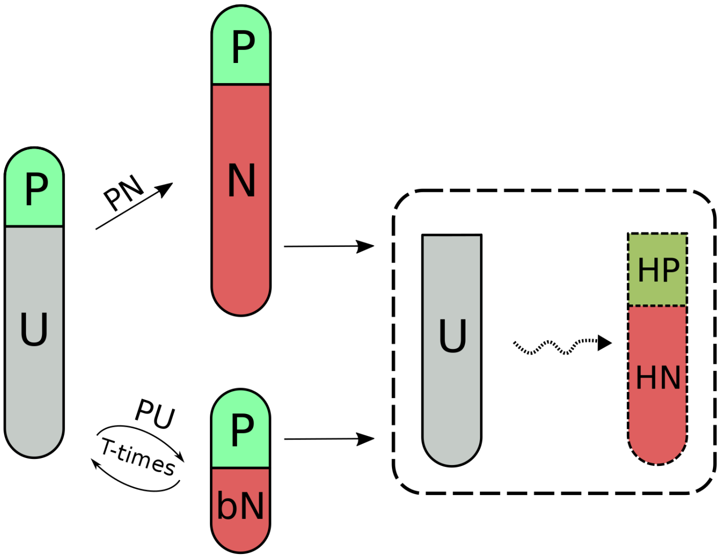


**Figure S2. Positive-Unlabeled bagging procedure and Positive-Negative learning difference.** Positive-unlabeled data in case of PN-learning is treated as positive and negative classes. For PU-bagging procedure, only a small part of unlabeled instances are treated as negatives, leading to significant decrease in false negative rate. Finally, both approaches attempt to identify hidden positives and hidden negatives in the unlabeled set.

P – positive training instances,

U – unlabeled instances,

bN – bootstrap negative instances,

N – instances treated as negative in PN-learning,
HP – hidden positive instances,
HN – hidden negative instances.

Hyperparameter is a parameter whose value is not explicitly learnt and needs to be tuned before the learning process (i.e. learning rate, tree depth etc.). Different classification algorithms (LR, SVM, AB, RF, DT) require different hyperparameters (**Table S2**). Process of finding the optimal set of hyperparameters that yields an optimal model is called hyperparameter optimization (tuning). We used grid search as the optimization technique.

Grid search is a brute-force algorithm that looks for the best subset of parameters through the whole parameters space (grid of hyperparameters). It iteratively takes each subset, trains the model with selected hyperparameters, evaluates the result and outputs the settings that achieved the highest score. Since PU data lacks negative examples, we used *PU*-score as a score function metric [11, 12]. As an evaluation technique we used 3-fold cross-validation: original sample was partitioned into 3 equal subsamples, one of them became test data and remaining ones used for training, cross-validation process was then repeated 3 times. Each subsample becomes test data only once. As a result, all three results obtained using testing data were averaged and final estimation was used to evaluate model performance during grid search. 3-fold cross-validation is then repeated for each subsample of hyperparameters.

Additionally, PU bagging has some specific parameters that need to be tuned. Firstly, number of bootstrap iterations in the bagging procedure, which affects redundancy of a final prediction. Mordelet et al. [10] observed little improvement in performance in bagging results after 100 bootstrap iterations. Therefore, we set 100 as a default number of iterations. Also, sampling coefficient could be used for modifying sampling procedure and number of unlabeled bootstrapped instances for training (*bN*, **Figure S2**). We set this coefficient to 1.0, resulting in the number of *bN* to be equal to the number of *P* (**Figure S2**).

After training and tuning, each ML algorithm is used to estimate the probability of *U* instances to belong to the Positive class. All the steps are repeated *T* times. For a given gene, probability of belonging to the Positive class is estimated as a mean of all probabilities for this gene obtained in the bagging procedure. All the steps are repeated for each classification algorithm.

<u>*Combining results from multiple ML algorithms*</u>

For finding an optimal combination of individual ML approaches, first, a formal metric for performance quality should be introduced. One of the ways to evaluate classification quality is to use a set of previously untouched instances of known nature.

GPrior uses a second set of true positive genes called algorithm evaluation set (AES) for the estimation of prediction error rates.

If a positive example was falsely classified as negative, it is called *FN* error. Overwise, when a negative example was falsely classified as positive, it is called *FP* error. Conventionally, *F1*-score metric is used in PN-learning as one of the measures of classification quality. Unfortunately, *F1*-score could not be obtained from the PU-data because it requires knowledge of *FP*. Since “true negative” data points that falsely were classified as positives could not be identified in PU-data, any metric depending on *FP* could not be applied for quality evaluation.

In case of gene prioritization problem - inability to confidently identify true negative genes makes it impossible to reliably calculate *FP*. Therefore, instead of *F1*-score, a similarly behaving metric, suitable for PU-learning, called *PU*-score was introduced [11, 12]:

$$PU_{Score} = \frac{p \times r}{Pr(y = 1)} = \frac{r^2}{Pr(\hat{y} = 1)}$$

where $p$ is precision,
$r$ is recall,
$Pr(y = 1)$ is the fraction of known positive labels in the predicted set,
$Pr(\hat{y} = 1)$ is the fraction of positive predictions made by the classifier.

Importantly, in terms of PU-learning, both recall and $Pr(\hat{y} = 1)$ can be calculated using only positive examples and *PU*-score could be used to evaluate performance of each ML approach and their possible combinations. *PU*-score metric is not limited to PU-learning and could as well be applied to conventional PN-learning methods.

Approach for finding the optimal combination of predictions is summarized in **Algorithm 1**. First, we define a set $B = \{b_1, \dots, b_n\}$, where $b_i$ is a final prediction of *i*-th classifier and *n* is the total number of used PU bagging classifiers (**Algorithm 1.1**). From *B* we can create a power set $W = P(B) \backslash \emptyset$, size of *N*, that stores all possible subsets of *B*, excluding empty set ∅ (**Algorithm 1.2**). Total number of subsets *N* equals to:

$$N = \sum_{k=1}^{n} \binom{n}{k}$$

For each subset of *W* we calculated average prediction *P* and corresponding *PU*-score **(Algorithm 1.3-6)**. *PU*-score is calculated using algorithm evaluation set (AES) to ensure independence of training and algorithm selection steps.

Conclusively, the best prediction that has the highest *PU*-score on AES is returned, resulting in a vector of probabilities corresponding to the genes in the input matrix **(Algorithm 1.7-12)**.

**Algorithm 1. Optimal combination of predictions**

(1) $B \leftarrow \{b_1, b_2 \ldots b_n\}$

(2) $W \leftarrow \mathcal{P}(B) \setminus \emptyset$

(3) $\mathrm{pu}_{max} \leftarrow 0$

(4) **for** each subset $w_i \in W$ **do**

(5) $\quad \mathrm{P} \leftarrow \textsc{average}(w_i)$

(6) $\quad \mathrm{pu} \leftarrow \textsc{pu-score}(\mathrm{P})$

(7) $\quad\quad$ **if** $pu > pu_{max}$ **do**

(8) $\quad\quad\quad \mathrm{p}_{max} \leftarrow \mathrm{pu}$

(9) $\quad\quad\quad \mathrm{opt} \leftarrow \mathrm{P}$

(10) **end for**

(11) **return** opt

**Algorithm 1. Description of the algorithm for finding optimal combination of the predictions.**
First, set of predictions *B* (1) is used for generating power set *W* (2), containing all of the possible combinations of predictions excluding empty set. We set maximum *PU*-score to zero (3) and start to iterate through the power set *W* (4). Each subset $w_i$ is then averaged (5) and evaluated by *PU*-score (6). If the obtained value is the highest so far (7), then maximum *PU*-score variable refreshes and this combination becomes the optimal (9). It is a brute-force algorithm, meaning that it takes into consideration all of the possible combinations, without additional optimization. Thus, it iterates through all subsets of $W$ and only then terminates iteration (10) and returns the optimal result (11).

*Prediction Quality Evaluation*

PU-learning approach imposes limitations on the usage of conventional predictor quality metrics – ROC, precision-recall analysis, etc. We used specific ways of prediction quality evaluation, not requiring knowledge of true negative instances.

1. *Gain/Lift charts*

As a result of prediction, a model returns a ranked list of genes of length *N*. Assuming that validation set (**Figure S1**) consists of *n* genes, most informative model should place all *n* genes at the top *n* positions of the list. Otherwise, if the prioritization is random the probability to observe these *n* genes at the top of the list is *n/N*.

The *lift* in this case is the number of genes detected by a model above a completely random selection of genes. The *gain* is the percentage of all positive examples from the validation set that have been found in the top subset of predictions (top 1%, 5%, 15%, 25% etc.).

Depending on the aim of analysis, it would make sense to analyze only the top percentage of the Gain/Lift chart, since for gene prioritization purposes overall quality of the classification is not so valuable as at the top of the sorted list. We used 1%, 5%, 15%, 25% thresholds for further model comparisons.

2. *Permutation rank test (Enrichment)*

Let's assume that we have a validation set of genes VS (**Figure S1**), each gene has its own rank in the overall prediction list *L* sorted by probability {*1,…, N*}. We expect that disease relevant genes will be ranked within the top percentiles (have very low ranks). Therefore, the extent to which a result could be obtained by chance could be tested by sampling *n* random genes from *L*, calculate the sum of their ranks, repeat this step *k* times (*k*=100,000 - 100,000,000) and calculate an empirical p-value. In this case p-value is the number of random sets which show the sum of ranks less or equal to the sum of ranks of VS divided by the number of samplings - *k*.

3. *PU-chart*

To estimate *PU*-score, first, the fraction of positive predictions made by the classifier should be estimated. This could be done by taking the sum of TP and FP divided by total number of instances. In this case we need to approximate the optimal decision boundary. Below this threshold will be all negative predictions, above all positive predictions. *PU*-chart visualizes the approximation process, showing the optimal threshold that maximizes *PU*-score. On *y*-axis there is *PU*-score, on *x*-axis – fraction of positive predictions made by the classifier.

4. *Statistical hypothesis testing methods*

We used a one-sided Mann–Whitney U-test to evaluate whether classification scores returned by the models for validation set are significantly higher than the ones given to remaining genes from unlabeled set.

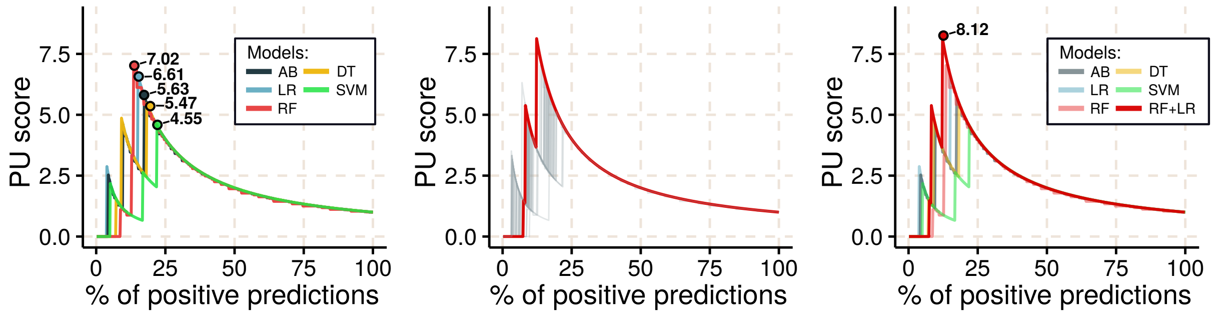


**Figure S3. Optimal combination of the predictions**. **(A)** Quality of each model is evaluated using PU-score metric; **(B)** All possible combinations of 5 models are evaluated and the one with the highest PU-score is selected for predictions. An independent set of genes – algorithm evaluation gene set is used to evaluate each combination; **(C)** Optimal combination of predictive models performs better than individual PU-models (Optimal combination - RF+LR).

### GPrior Performance Evaluation

#### *Wisconsin breast cancer dataset*

We used Wisconsin breast cancer data set in order to introduce the PU-learning concept and compare PU and PN approaches in terms of performance with relation to percentage of known positives (KP). It was created by Dr. William H. Wolberg, physician at the University of Wisconsin Hospital at Madison, Wisconsin, USA. It consists of 32 attributes (30 features) and 569 instances. Features are computed from a digitized images aspiration of a breast mass. Most of them describe characteristics of the cell nuclei. Importantly, tumors are classified into malignant and non-malignant and the true classifications could be used for evaluation ML classification performance.

Source: https://archive.ics.uci.edu/ml/datasets/Breast+Cancer+Wisconsin+(Diagnostic)

#### *GPrior performance comparisons*

GPrior estimates *PU*-scores from each of 5 algorithms used and probes all possible combinations of predictions to select the one with the largest *PU*-score estimated using algorithm evaluation set (**Figure S3**).

We used PCA to highlight malignant and non-malignant tumors in the benchmarking dataset (**Figure S4A**). It could be seen that there is no clear clustering of the data, rather the differences between classes are very minor. As a result, GPrior (and

PU-based learning methods), are able to detect similarity to the positive instances much more efficiently than conventional PN-based learning algorithm (**Figure S4B,C**).

Available true positive and true negative classifications in the Wisconsin breast cancer dataset allow to evaluate performance of the methods with both conventional *F1*-score and *PU*-score (**Figure S4D,E**). GPrior, as an ensemble PU-learning performs better than individual PU-learning approach (bagging SVM), which is a part of GPrior and conventional PN-learning (biased SVM) in case of small fraction of true instances being known.

It is only with more than 20% of known true positive instances used for training when PN-learning starts to perform nearly equally well. Knowing 20% of genes involved in a complex trait is yet an unsolved case for the majority of phenotypes.

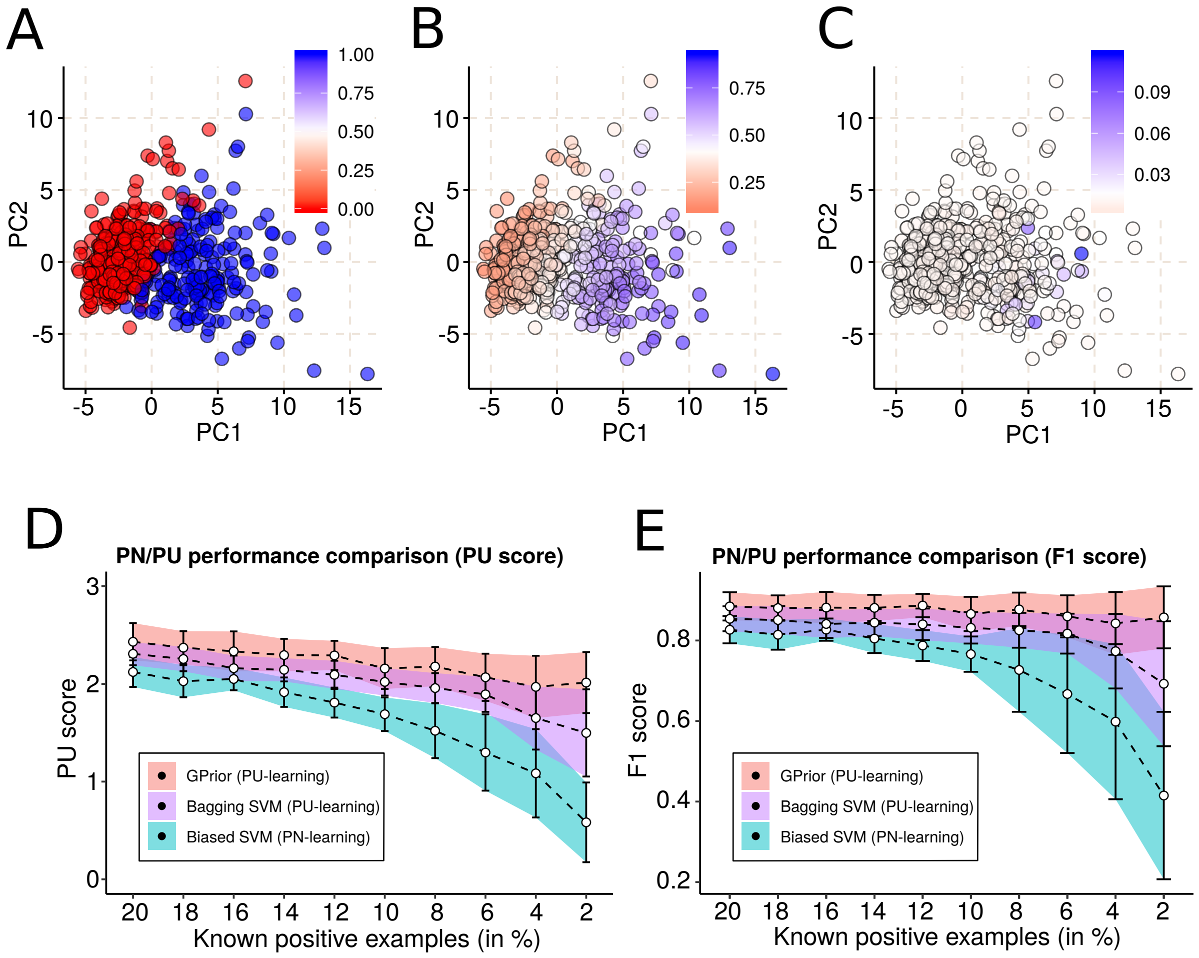


**Figure S4. Evaluation of PN and PU learning methods performance using benchmark dataset.**

**(A)** PCA of the original data, showing non-malignant (true negative, red) and malignant (true positive, blue) tumors in Wisconsin Breast Cancer data; **(B)** PCA of the original data colored with respect to the probability of an instance coming from true positive (blue) class. Probabilities estimated using GPrior; **(C)** PCA of the original data colored with respect to the probability of an instance coming from true positive (blue) class. Probabilities estimated using biased SVM (PN-learning); **(D)** Comparison of algorithm performances for different fraction of true positive instances used for training using *PU*-score; **(E)** Comparison of algorithm performances for different fraction of true positive instances used for training using *F1*-score.

Changing the fraction of known positives not just provides more information to train the model, but additionally, creates a different fraction of hidden positives in the unlabeled set. To highlight only methodological difference effects between PN and PU-learning algorithms we fixed the fraction of hidden positives in the unlabeled set.

First, the true positive set (N=212) was separated into two parts – one of them was fixed and always played a role of hidden positives in the unlabeled set (N=119) and the other was used to randomly draw appropriate number of true positive instances for training (N=93) (**Figure S5A**). Such approach results in a subset of the original benchmarking dataset with fixed fractions of hidden positives and negatives in the unlabeled set (**Figure S5B**).

Finally, upon examining different fractions of known positive instances in the dataset, we confirmed superior performance of PU-learning and specifically, GPrior (**Figure S5C**).

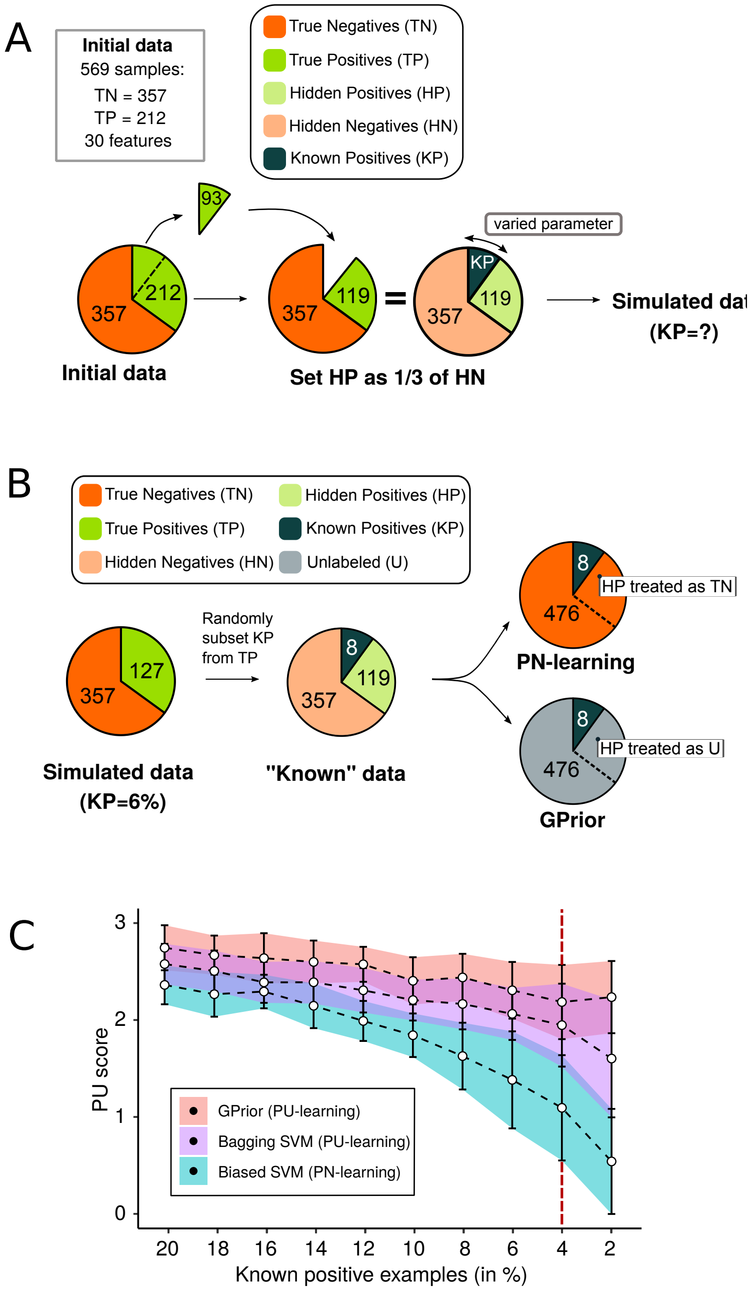


**Figure S5. Benchmark dataset analysis with fixed fraction of hidden positive instances in the unlabeled set.**

**(A)** Scheme for construction of the benchmark data subset with fixed fraction of hidden positive instances in the unlabeled set; **(B)** Training scheme for PN and PU-learning; **(C)** *PU*-score based comparison of performance evaluation for biased SVM, bagging SVM and GPrior.

### Case studies

#### *Educational attainment*

One of the challenges of gene prioritization problem is a quality evaluation. We used several approaches to solve this issue – PU-score, gain/lift charts, but so far we have not addressed the question of specificity of the prediction. We tested whether GPrior returns trait-specific prioritization by conducting a control experiment with two highly distinct phenotypes (IBD and EA). We hypothesized that usage of irrelevant training set should return no notable prioritization of true disease genes.

Summary statistics for EA was taken from Lee et al [13]. This study used one of the largest sample sizes for EA - 1,131,881 individuals. Pre-processing resulted in the gene-based data matrix with 10,739 gene candidates found in loci with original p-value < $10^{-6}$.

Training and validation sets were constructed using GWAS Catalog (EFO_0004784). We kept only regions mapped to a single gene and sorted them based on p-value. All genes with associated p-value in range $2x10^{-95}$ to $10^{-13}$ were used for training. First 50 of them became true gene set, remaining 69 became algorithm evaluation set. Genes (n=381) with p-value from $10^{-13}$ to $10^{-6}$ became the validation set (**Sup. Table S6**).

To eliminate potential bias in the size of training sets for the two phenotypes, we used for GPrior training only 18 genes (12 for ML training and 6 for algorithm evaluation) from the IBD training gene set that were also found in EA GWAS loci with p-value < $10^{-6}$. As validation sets we used original validation sets for IBD and EA.

We observed trait-specific enrichment signal only in case of using disease-relevant training set, confirming trait-specific nature of GPrior prioritizations and, additionally, showing the suitability of approach with usage of independent validation gene set for quality evaluation (**Figure S6**).

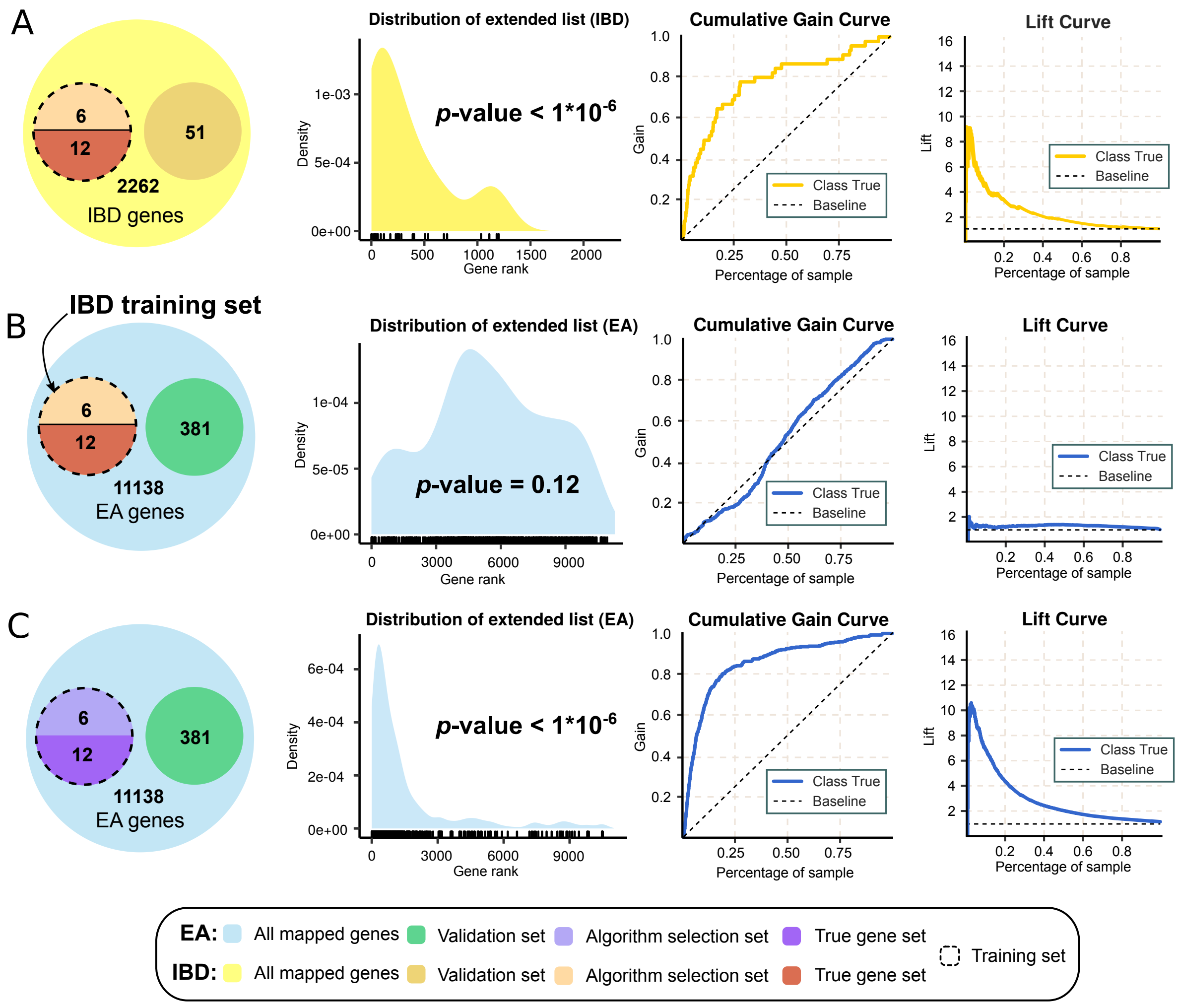


**Fig. S6. Control experiment for IBD genes prioritization using EA GWAS.**
(A) Dataset breakdown and fixation of contamination by setting a single number of HP instances to be used for all simulations; (B) Scheme for the simulation with fixed contamination; (C) Performance of PU and PN learning approaches with respect to a fraction of known positive data points.

Next, we applied GPrior to EA data, using a full-size training list (**Figure S7, Table S7, S8**) and confirmed the best quality of GPrior predictions out of all tools used for comparison.

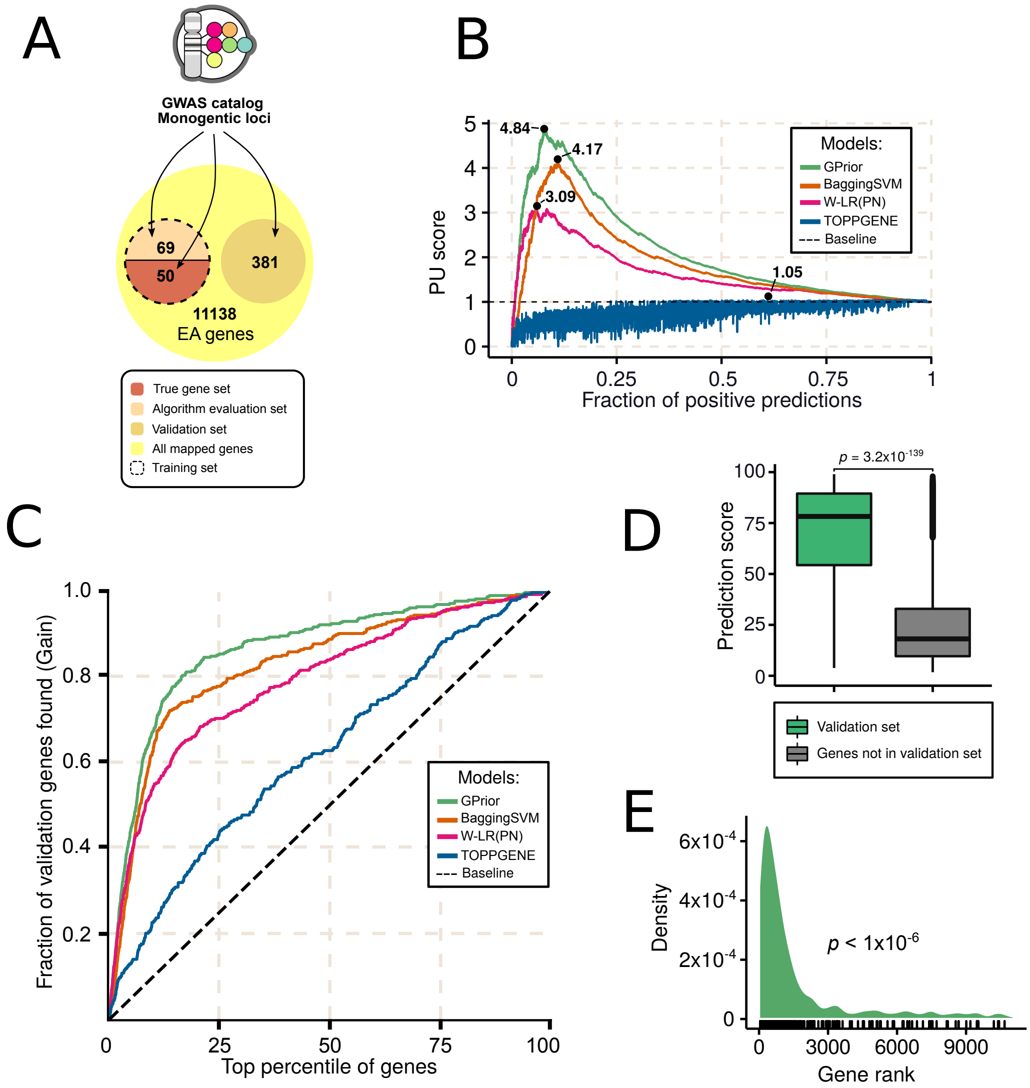


**Figure S7. GPrior results for educational attainment GWAS.**
**(A)** Scheme for selection of training, algorithm evaluation and validation gene sets; **(B)** Quality comparison for GPrior, Bagging SVM and conventional PN-learning with linear regression; **(C)** Cumulative gains curve shows better prioritization of true genes using GPrior in comparison with other methods, including TOPPGENE; **(D)** GPrior prediction score for genes found in genome-wide significant loci. True genes from the independent validation gene set receive significantly higher scores than genes found within the same locus but not implicated in the disease; **(E)** Enrichment of true genes from independent validation gene set among top predictions from GPrior.

Next, we sought to assess the contribution of the variant-to-gene mapping into the prediction quality. Functionality for gene prioritization is still in early development in POSTGAP, yet, it reports a "variant-to-gene" mapping quality score based on 7 features. Most of them are variant functional annotations, indicative of whether an SNP is found within, near or affects regulation of a specific gene (VEP, Nearest, GTEx, Fantom5, DHS, PCHiC, Regulome) [1]. We used maximal variant-to-gene mapping scores for each gene to construct a ranked list of genes and computed the *PU*-score for such prioritization scheme, using 381 genes from validation set (PU = 3.82). Importantly, such variant-to-gene score is not directly related to the question of whether the gene is disease-relevant.

We used exactly the same 7 features to run analysis with GPrior, yielding (PU = 4.1). GPrior could use a significantly greater number of meaningful features, therefore, upon using the whole data matrix with relevant features, performance increased even more (PU = 4.84, **Figure S7B, Figure S8**).

We used a random forest model to analyze feature importance in the training data to highlight contribution of variant-to-gene mapping quality. Notably, variant functional annotations and location of the nearest gene to a particular variant are of high importance (**Figure S8**). Therefore, we expect that careful fine-mapping should significantly improve the quality of prioritizations from any model. Additionally, GPrior uses a non-finite set of features and would benefit from usage of disease-specific features.

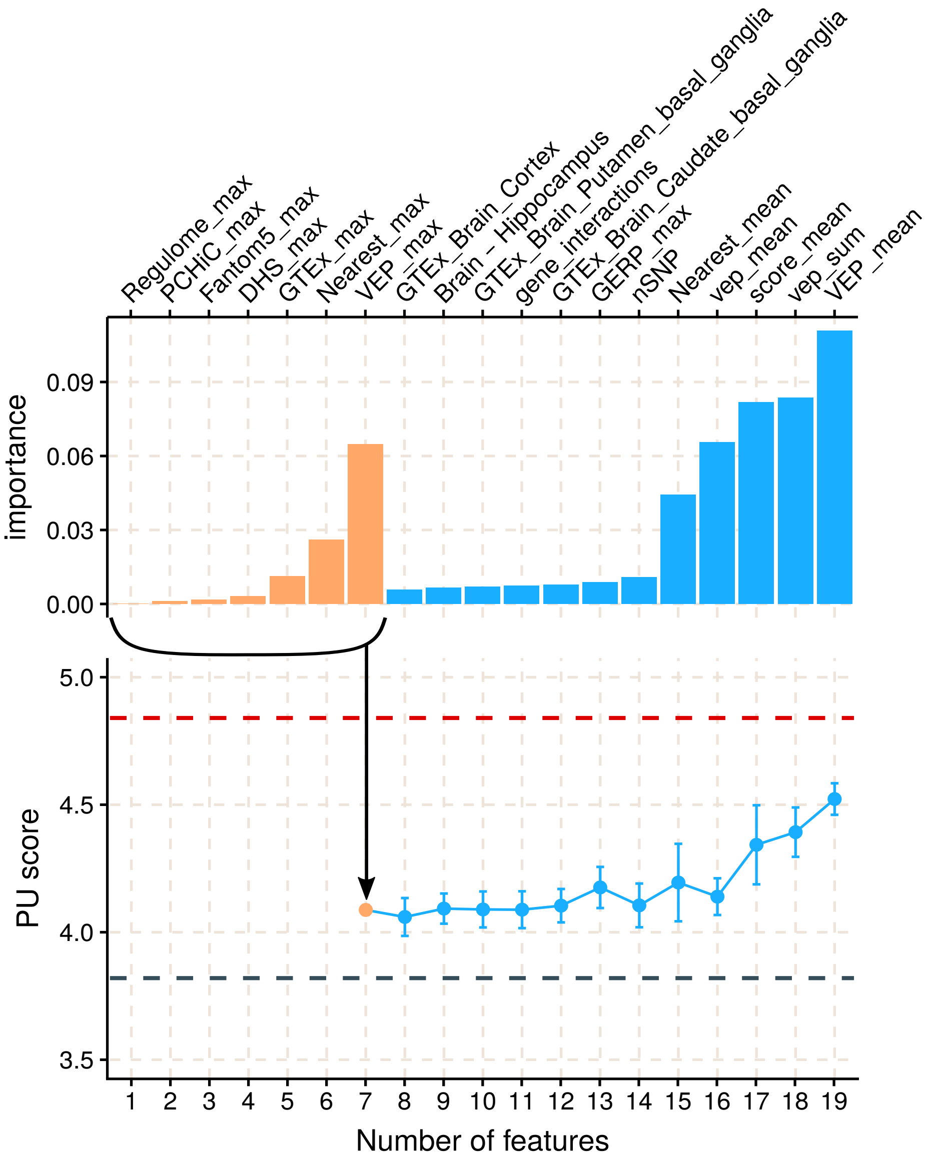


**Figure S8. Feature importance and variant-to-gene mapping contribution.**
Though not intended for gene prioritization, POSTGAP that we used for variant-to-gene mapping and assembly of the candidate gene list for prioritization, reports a variant-to-gene mapping score that could be used to rank genes. This score is estimated as a sum of 7 features (top panel; orange). PU-score for a classifier based on this custom score is represented by the dashed grey line. If GPrior is used on exactly the same input parameters and restricted to the same 7 features we get ~11% increase in PU-score. Finally, GPrior takes advantage of the large number of additional features that could not be introduced to the POSTGAP model, which results in further improvement of the PU-score (blue). Importantly, feature importance highlights greater value of the variant-level features for gene prioritization.

### Comparison with MAGMA

So far, we have compared GPrior with other ML-algorithms (BaggingSVM; W-LR) and gene prioritization techniques (TOPPGENE). Altogether, GPrior demonstrated best performance in comparison with all of them. In this series of experiments we tried to compare GPrior results with an already existing similar gene prioritization method (MAGMA) that takes GWAS summary statistics as an input [14]. We ran MAGMA with default parameters according to the manual.

For each phenotype MAGMA produced a smaller number of gene candidates than were used as an input data matrix for GPrior. It is important for the prioritization tool to produce significant enrichment of the disease genes at the very top of the ranked list of candidates. We extracted only top *n* genes from GPrior results, where *n* is the size of MAGMA output. For example, for SCZ summary statistics from Pardiñas et al. [15], MAGMA produced a gene set size of 674. GPrior, on the other hand, produced 3903 genes, including training set. To make the comparison fair, we excluded all of the genes used for training and took only the top 674 from the sorted list for GPrior. Further, we computed an enrichment of the validation set genes in at the top of the predictions list.

For each phenotype we used the same validation sets that were used at the dedicated case studies. The only difference is that validation sets contained only genes found in both MAGMA and GPrior prioritizations. Enrichment of the validation genes was computed with permutation test (**Figure S9**).

For all of the traits GPrior demonstrated enrichment of top ranked predictions for validation set for all phenotypes – IBD (P=0.019), EA (P=9e-03), CAD (P=3e-3) and schizophrenia (P=7e-4). MAGMA produced significantly enriched predictions for CAD (P=9e-03).

Conclusively, GPrior demonstrated best performance out of all evaluated approaches for gene prioritization in multiple settings and for various phenotypes.

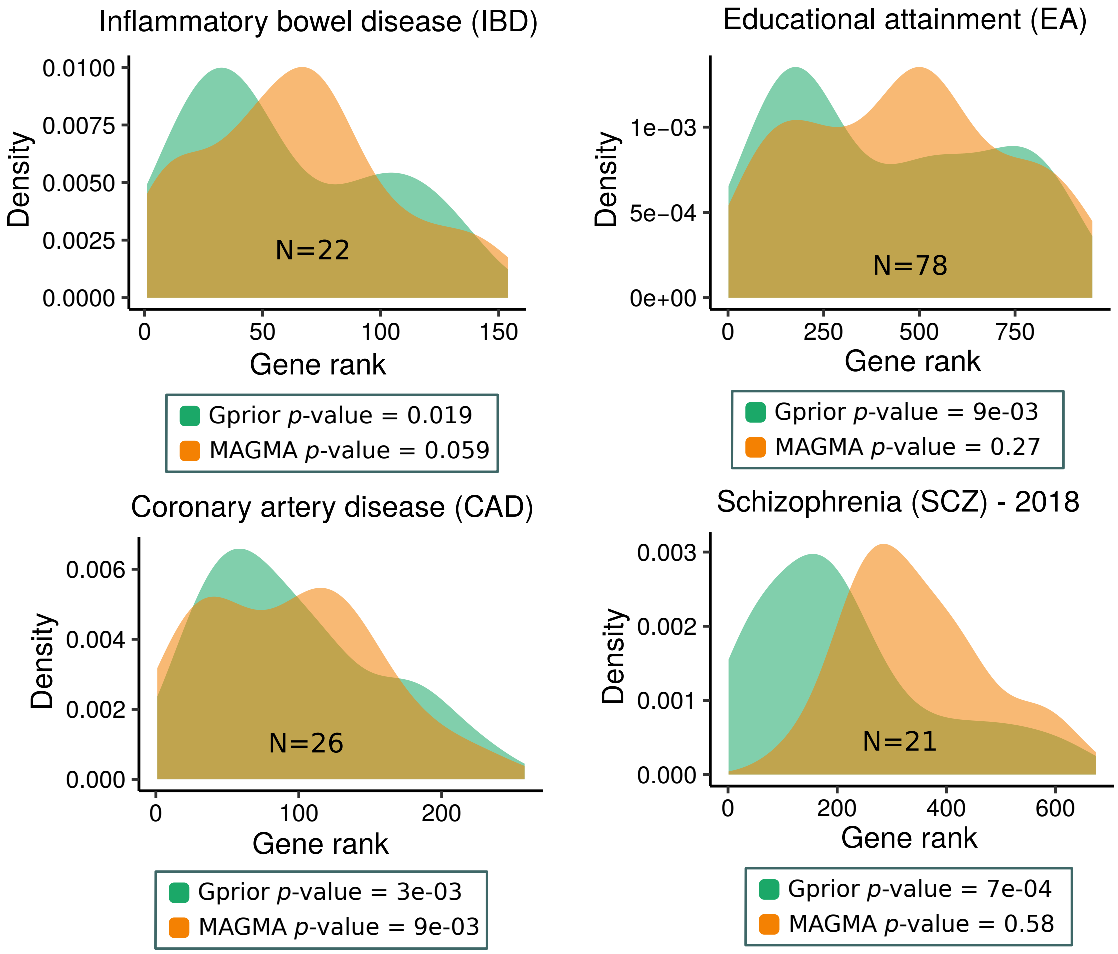


**Figure S9. Comparison of GPrior and MAGMA prioritizations.**
